# Bone morphogenetic protein 9 induces osteogenic differentiation of germ cell 1 spermatogonial cells

**DOI:** 10.1101/823435

**Authors:** Jiye Zhang, Bangfa Xu, Xinping Chen, Liqiang Zhao, Pei Zhang, Fei Wang, Xiaojuan Li, Meiling Wang, Weihua Xu, Wenwen Zhang, Shengmiao Fu

## Abstract

Germ cell 1 spermatogonial (GC-1spg) cells are multipotent progenitor cells. We previously confirmed that bone morphogenetic protein (BMP) 9 is among the most osteogenic BMPs. However, whether GC-1spg cells are driven toward osteogenic differentiation under proper stimuli is uncertain. Additionally, the molecular mechanism of BMP9 remains unclear. In the present study, we aimed to determine whether BMP9 can induce osteogenic differentiation of GC-1spg cells. Recombinant adenoviruses were generated by the AdEasy system to regulate the BMP9 expression in GC-1spg cells. We identified osteogenic markers by real-time PCR and staining techniques *in vitro*. Ectopic ossification assays and histological analysis were also performed to verify the *in vivo* activity of BMP9. Finally, potential signaling pathways of BMP9 were assessed by transcriptome sequencing and KEGG enrichment analysis. Using recombinant adenoviruses, we demonstrate that BMP9 upregulates osteogenic markers including Runx2, osteocalcin, osteopontin, and Sox9. BMP9 also activates alkaline phosphatase activity and calcium deposition in GC-1spg cells. *In vivo* results show that BMP9 overexpression in GC-1spg cells promotes ectopic bone formation and chondrogenesis. In addition, RNA-sequencing and KEGG pathway analysis demonstrate that several signaling pathways are involved in BMP9-mediated osteogenesis. GC-1spg cells not only maintain spermatogenesis but also retain the ability to form bone tissue. Therefore, BMP9 activity in GC-1spg cells may help identify signaling pathways implicated in bone formation and could be of use in regenerative medicine.

## Introduction

Germ cell 1 spermatogonial (GC-1spg) cells are mouse spermatogonia that are immortalized by the SV40 large T antigen [1]. These cells can differentiate into diverse cell types or undergo self-renewal, depending on different factors [2,3,4]. GC-1spg cells are largely involved in spermatogenesis and have mesenchymal stem cell (MSC) differentiation potential [5,6,7,8]. In other words, GC-1spg cells can differentiate into different cell types, including those of the osteogenic, chondrogenic, or adipogenic lineages [9,10,11,12]. However, several molecular events [13,14] and factors are involved in osteogenic differentiation of MSCs. Importantly, bone morphogenetic proteins (BMPs) are essential for this process [15,16,17].

BMPs, members of the TGF-β superfamily, are required for bone formation, stem cell differentiation, and male reproduction [15,16,18,19,20]. Thus far, more than 15 different BMPs have been identified. Our previous work showed that BMP9 is the most osteogenic BMP [21,22,23,24]. BMP9, also known as growth differentiation factor 2 (GDF-2)[25], was originally isolated from the fetal mouse liver [25,26]. Bone formation involves BMP9 acting through the Notch signaling pathway [14,24,27]. Several metabolic processes are regulated by BMP9, including glucose and lipid metabolism [28,29], iron metabolism [30], endothelial function, and angiogenesis [31,32,33]. Accordingly, abnormal BMP9 expression may be involved in various diseases [29,34,35,36].

Studies confirm that GC-1spg cells have MSC characteristics [4,7,12,37], however, few reports have focused on the BMP9-mediated osteogenic effect on GC-1spg cells. Indeed, MSCs can be isolated from the testis [5,6]. We analyzed how GC-1spg cells react to BMP9 and demonstrate that BMP9 can induce bone formation in GC-1spg cells both *in vitro* and *in vivo* [10,17,38]. In addition, we analyzed the GC-1spg cell transcriptome after BMP9 treatment [27] and investigated potential BMP9 signaling pathways by KEGG enrichment analysis.

## Materials and Methods

### Cell culture and chemicals

GC-1spg and HEK-293 cells were purchased from the American Type Culture Collection (ATCC). Cells were cultured in complete Dulbecco’s Modified Eagle Medium (DMEM, HyClone, Logan, UT, USA) supplemented with 10% fetal bovine serum (FBS, HyClone), 100 U/ml penicillin, and 100 g/ml streptomycin. Cultured cells were incubated at 37 °C in 5% CO_2_[1,39,40,41]. Unless otherwise specified, all chemicals were purchased from Sigma-Aldrich (St. Louis, MO, USA) or Thermo Fisher Scientific (Waltham, MA, USA).

### Recombinant adenovirus construction

To regulate BMP9 expression, recombinant adenoviruses were generated by the AdEasy system [22,39,42,43]. Human BMP9 DNA was amplified by high-fidelity PCR as follows: 96°C for 45 s, followed by 18 cycles at 92 °C for 20 s, 55 °C for 30 s, and 70 °C for 45 s, and a final 70 °C incubation for 5 min. The amplified sequence was cloned into the adenoviral shuttle vector, which was transformed into HEK-293 cells to generate recombinant adenoviruses [39]. The resultant adenoviruses (Ad-BMP9) also expressed green fluorescent protein (GFP). Control adenoviruses expressing only GFP (Ad-GFP) were also constructed [17,38].

### RNA isolation, reverse transcription, and real-time PCR

Total RNA from GC-1spg cells was extracted using TRIZOL reagent (Aidlab, Beijing, China). Random hexamers and M-MuLV Reverse Transcriptase (VAZYME, Nanjing, China) were used for reverse transcription. Reverse transcription reactions were carried out under the following conditions: 25 °C for 5 min, 50 °C for 15 min, 85 °C for 5 min, and 4 °C for 10 min. The cDNA was further diluted by 10-fold and used as real-time PCR templates. All PCR primers were designed using Primer3web software (Table 1) [17,38,44,45]. Target sequences were incubated as follows: 50 °C for 2 min and 95 °C for 10 min, followed by 40 cycles at 95°C for 30 s and 60 °C for 30 s. Real-time PCR assays were performed on an ABI QuantStudio 6 instrument. The reagents included diluted cDNA (4 µL), forward and reverse primers (0.4 µL each), SYBR Green Master Mix (10 µL), 50× ROX Reference Dye 2 (0.4 µL), and H_2_O (4.8 µL). Relative gene expression data were analyzed by the 2^-ΔΔCt^ method [46,47]. Assays were performed with at least three independent biological replicates.

**Table 1.**
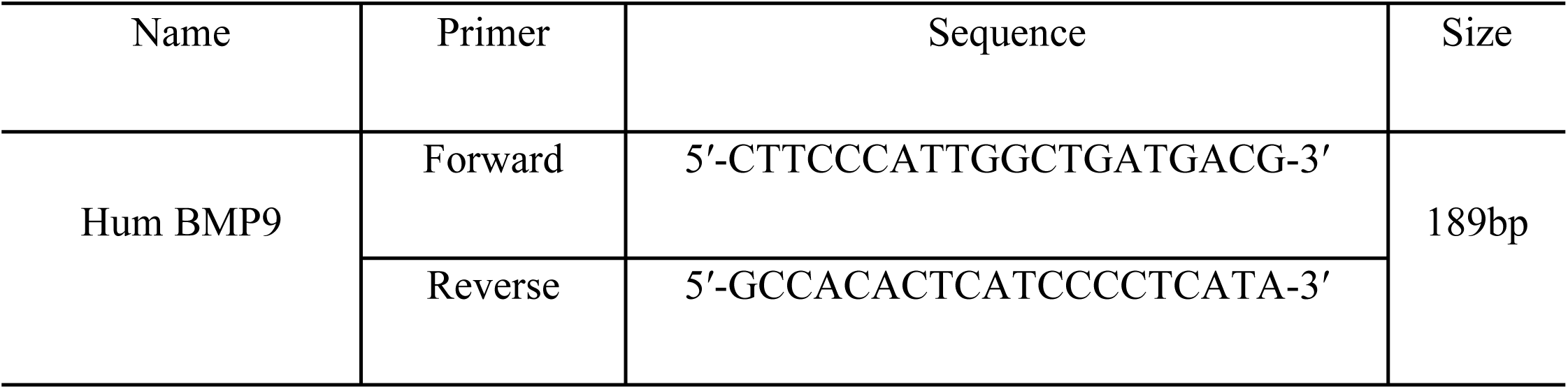
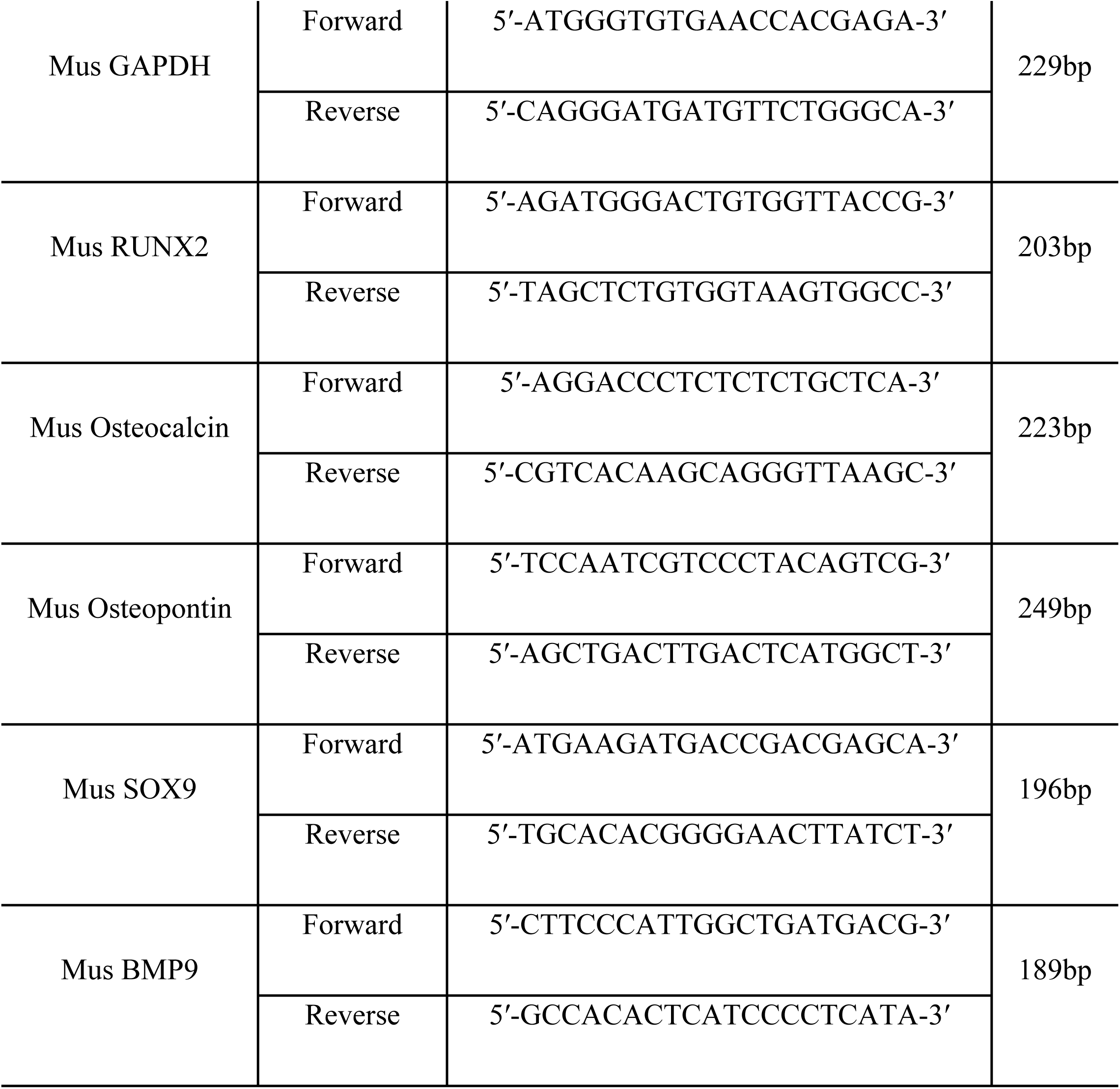
Real-time PCR primers.

### Alkaline phosphatase (ALP) activity assays

ALP activity was assayed by histochemical staining [17,38,48,49,50]. Log-phase GC-1spg cells were seeded in SlideFlasks (Thermo Fisher Scientific) and infected with Ad-BMP9 or Ad-GFP. At the indicated time points, ALP activity was assessed using a BCIP/NBT Chromogen assay kit (Solarbio, Beijing, China). Assays were repeated at least three times.

### Alizarin Red S staining

Alizarin Red S staining was used to identify mineralized matrix nodules or calcium precipitation [17,38,51]. GC-1spg cells were seeded in SlideFlasks, infected with Ad-BMP9 or Ad-GFP adenoviruses, and maintained in DMEM containing ascorbic acid (50 mg/mL) and β-glycerophosphate (10 mM) for 7 or 9 days [51]. Culture medium was then discarded, and the cells were carefully washed thrice with PBS. Cells were then fixed with 1% glutaraldehyde at room temperature for 10 min and rinsed thrice with PBS. Next, cells were incubated for 30 min at approximately 28 °C in 0.2% Alizarin Red S (Solarbio) and rinsed thrice with PBS. Calcium mineral deposits were observed with bright field microscopy.

### Cell implantation and ectopic ossification assay

Animal experiments were approved and supervised by the research ethics committee of Hainan General Hospital. Ectopic bone formation assays were performed as previously described [17,38,52,53,54]. First, sub-confluent GC-1spg cells were treated with Ad-BMP9 or Ad-GFP for 24 h. Cells were harvested (5×10^6^ cells per sample) and resuspended in 100 µL PBS. Transformed cells were subcutaneously injected into the flanks of 4-week-old male athymic nude mice (5 animals per group). After 4 weeks, mice were euthanized, and injection-site tissue was collected for further analysis. Characteristic masses were selected, fixed in 10% buffered formalin, and imaged using a micro-CT system (Skyscan 1076, Antwerp, Belgium). Tissue volume, bone volume, and bone surface area were measured for each tissue sample [55,56].

### Histological analysis

Collected tissue samples were fixed in 10% buffered formalin overnight and embedded in paraffin. Serial sections were stained with hematoxylin and eosin (H&E, Sigma-Aldrich), Masson’s trichrome (Sigma-Aldrich), and alcian blue (Sigma-Aldrich) [17,27,38,53].

### Signaling pathway analysis

Transcriptome sequencing and KEGG enrichment analysis were performed for BMP9 relevant signaling pathway analysis. GC-1spg cells were seeded in 100 mm culture dishes and infected with Ad-BMP9 or Ad-GFP. Total RNA was collected 48 h post-infection. At least 6×10^7^ cells were used for RNA extraction, and all RNA from one group was pooled into one sample. Whole RNA-seq libraries, including lncRNA, circRNA, mRNA, and miRNA, were prepared by Novogene Bioinformatics Technology (Tianjin, China) and sequenced with an Illumina HiSeq 2000/4000 platform (San Diego, CA, USA). Finally, potential BMP9 signaling pathways were identified using KOBAS (2.0) software [57,58], which was used to calculate transcriptome enrichment for mRNA, lncRNA, circRNA, and miRNA. BMP9-related KEGG pathways were identified as described[59,60,61].

### Statistical analysis

Data were analyzed by GraphPad Prism 7 and SAS 9. Significant differences between Ad-BMP9 and Ad-GFP groups were determined by Student’s *t*-tests. Hypergeometric P-values < 0.05 were calculated to identify significant associations in KEGG pathways.

## Results

### BMP9 upregulates osteogenic differentiation markers in vitro

We first tested our recombinant adenoviruses and observed that Ad-BMP9 upregulated BMP9 expression (Fig. 1A-B). GFP expression was also present in Ad-BMP9-infected cells (Fig. 1A-B). BMP9 is a potent osteogenesis inducer. Therefore, we next tested BMP9 function in GC-1spg cells. Several osteogenic differentiation markers, including Runx2, osteocalcin (OCN), osteopontin (OPN), and Sox9 [17,38,53], were upregulated by BMP9 (Fig. 1C). We then examined the early osteogenic markers ALP [17,62,63] and mineral node formation. ALP activity was analyzed by histochemical staining. Our results showed that BMP9 overexpression increased early osteogenesis differentiation, indicated by ALP activity in GC-1spg cells (Fig. 1D). Alizarin Red S staining was then used to identify mineral nodes. Similarly, BMP9 overexpression visibly increased mineral nodule formation *in vitro* (Fig. 1E). These results suggest that BMP9 regulates osteogenic differentiation of GC-1spg cells.

**Fig 1.**
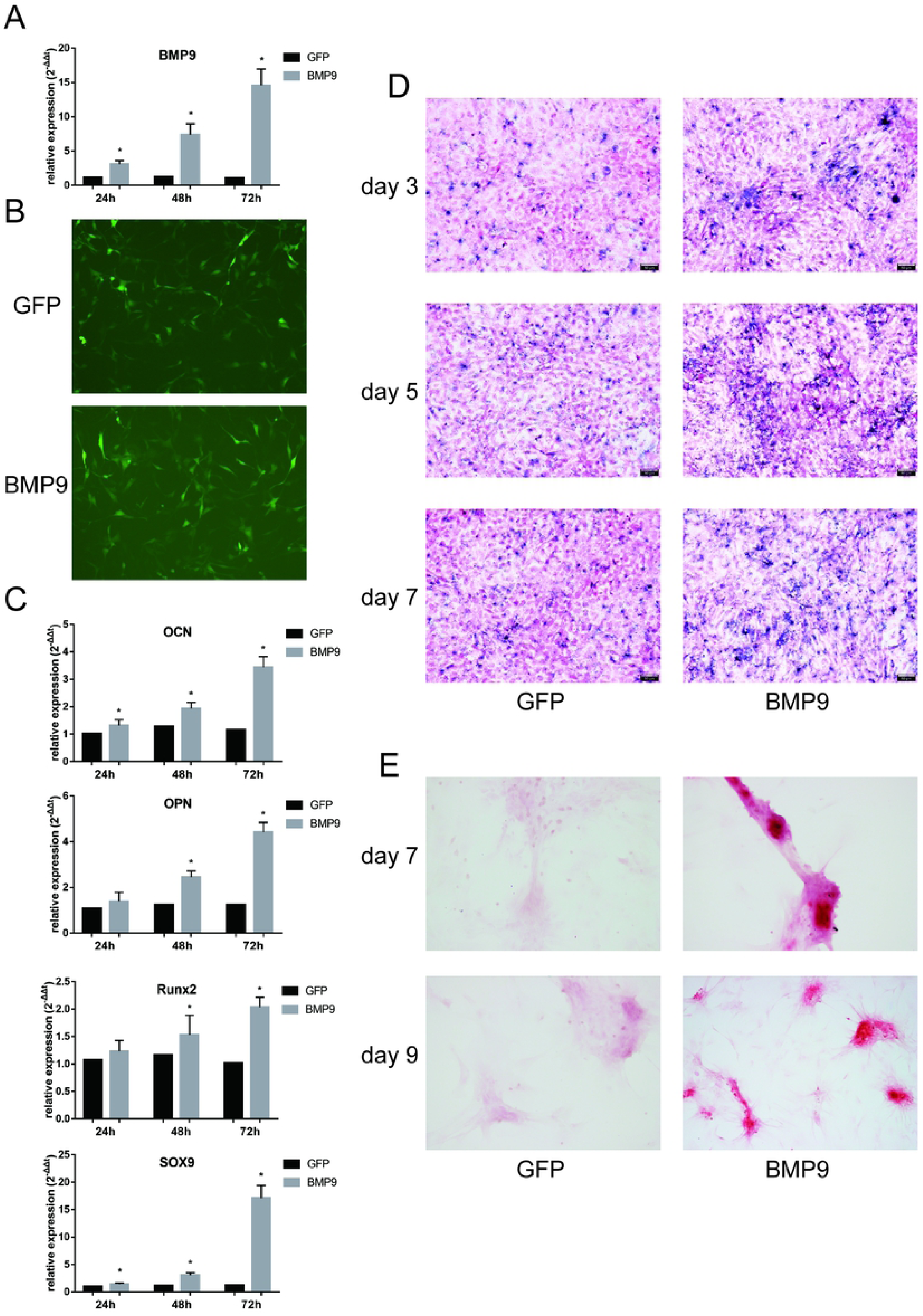
Exogenous BMP9 augments osteogenic differentiation marker expression and potentiates osteogenic differentiation [A & B] Recombinant Ad-BMP9 upregulates BMP9 expression in GC-1spg cells. The recombinant adenoviruses Ad-BMP9 and Ad-GFP effectively transduce GC-1spg cells. GFP was detected 24 h post-infection. Sub-confluent GC-1spg cells were treated with Ad-BMP9 or Ad-GFP for 24 h, 48 h, or 72 h. Total RNA was isolated and analyzed by real-time PCR. *P<0.05. [C] The recombinant adenovirus Ad-BMP9 mediates BMP9 overexpression in GC-1spg cells. At the indicated time points, treated cells were analyzed by real-time PCR. Osteogenic markers were upregulated by BMP9. *P<0.05. [D] Cells were analyzed by qualitative histochemical alkaline phosphatase (ALP) staining 3, 5, and 7 days after Ad-BMP9 treatment. BMP9 induces the early osteogenic marker ALP in GC-1spg cells (magnification, 200×). [E] Sub-confluent GC-1spg cells were treated with the indicated adenoviral vectors and cultured in mineralization medium for 7 or 9 days. The cells were fixed and stained with Alizarin Red S. Representative mineral nodule images are shown (magnification, 200×).

### BMP9 induces ectopic bone formation in vivo

To verify our *in vitro* results that show exogenous BMP9-induced GC-1spg cell osteogenesis, we performed ectopic ossification experiments. Using our previously established stem cell implantation assay [14,17,38,53], we detected injection-site masses in both BMP9 and GFP groups. Masses in the BMP9 group appeared slightly larger than in the GFP group. Micro-CT analysis also showed differences in tissue volume, bone volume, and bone surface area between Ad-BMP9 and Ad-GFP-injected tumors (Fig. 2 A-B).

**Fig 2.**
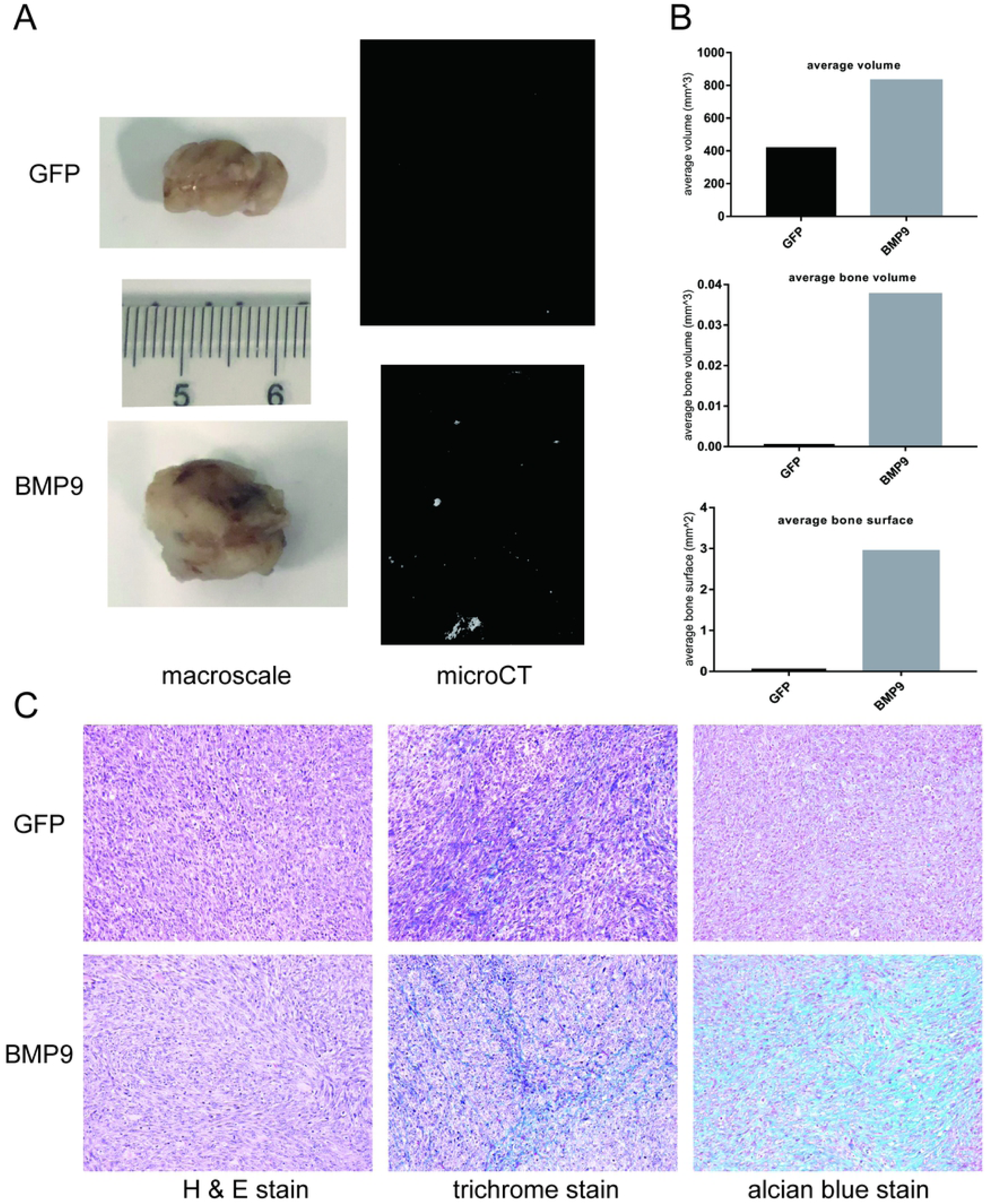
BMP9 induces mineralization and maturation of ectopic bone formations [A] Sub-confluent GC-1spg cells were treated with Ad-BMP9 or Ad-GFP for 16 h. Treated cells were collected for subcutaneous injections into athymic nude mice. Cells were allowed to grow for 4 weeks. Characteristic masses were retrieved and examined by micro-CT. [B] Injection-site products in the BMP9 group were larger than in the GFP group. Ossification structure volume and surface area of the masses in the BMP9 group were also higher than those of the GFP group. [C] Tumor samples were fixed, decalcified, embedded in paraffin, and stained with H&E, Masson’s trichrome, or alcian blue. Representative images are shown. The Ad-GFP group exhibited less mineralized, immature chondroid matrix compared to the Ad-BMP9 group.

Histologic analysis verified BMP9 function in GC-1spg cells. Significant differences were not observed between the two groups using H&E staining, but Masson’s trichrome staining revealed more mature cells with increased mineralization from exogenous BMP9 expression. Analogously, alcian blue staining also showed the accumulation of chondroid matrix in BMP9-induced GC-1spg cells (Fig. 2C). Taken together, our data demonstrate that BMP9 induces bone formation in GC-1spg cells.

### BMP9 activity is transduced by several cellular pathways

Several recent studies demonstrated that Notch signaling is required for BMP9-regulated bone formation in MSCs [14,27,48,64]. However, some studies also demonstrated that BMP9 can induce MSCs to differentiate into various cell types [22,65,66]. Moreover, aberrant BMP9 expression might be involved in certain diseases [20,29,34]. These studies suggest that several signaling pathways are involved in BMP9-regulated cell fate or proliferation, but the specific pathways are unclear. We analyzed total RNA, including lncRNA (Fig. 3A), miRNA (Fig. 3B), circRNA (Fig. 3C), and mRNA (Fig. 3D), by RNA-seq followed by KEGG pathway analysis. Our analysis showed that TGF-β, Notch, MAPK, and Ras signaling pathways mediate BMP9-induced differentiation. This result corroborates previous findings identifying the mechanism of BMP9-induced osteogenic differentiation [14,27,65,66,67,68]. Simultaneously, our results also revealed that insulin, PPAR, glutathione metabothyroid hormone, TNF, PI3K-Akt, and several cancer pathways are also involved in BMP9-mediated cellular differentiation. Thus, our findings suggest that BMP9 signaling may be targeted for the treatment for adipogenesis, diabetes, cancer, and certain chronic inflammatory diseases [17,28,29,69,70].

**Fig 3.**
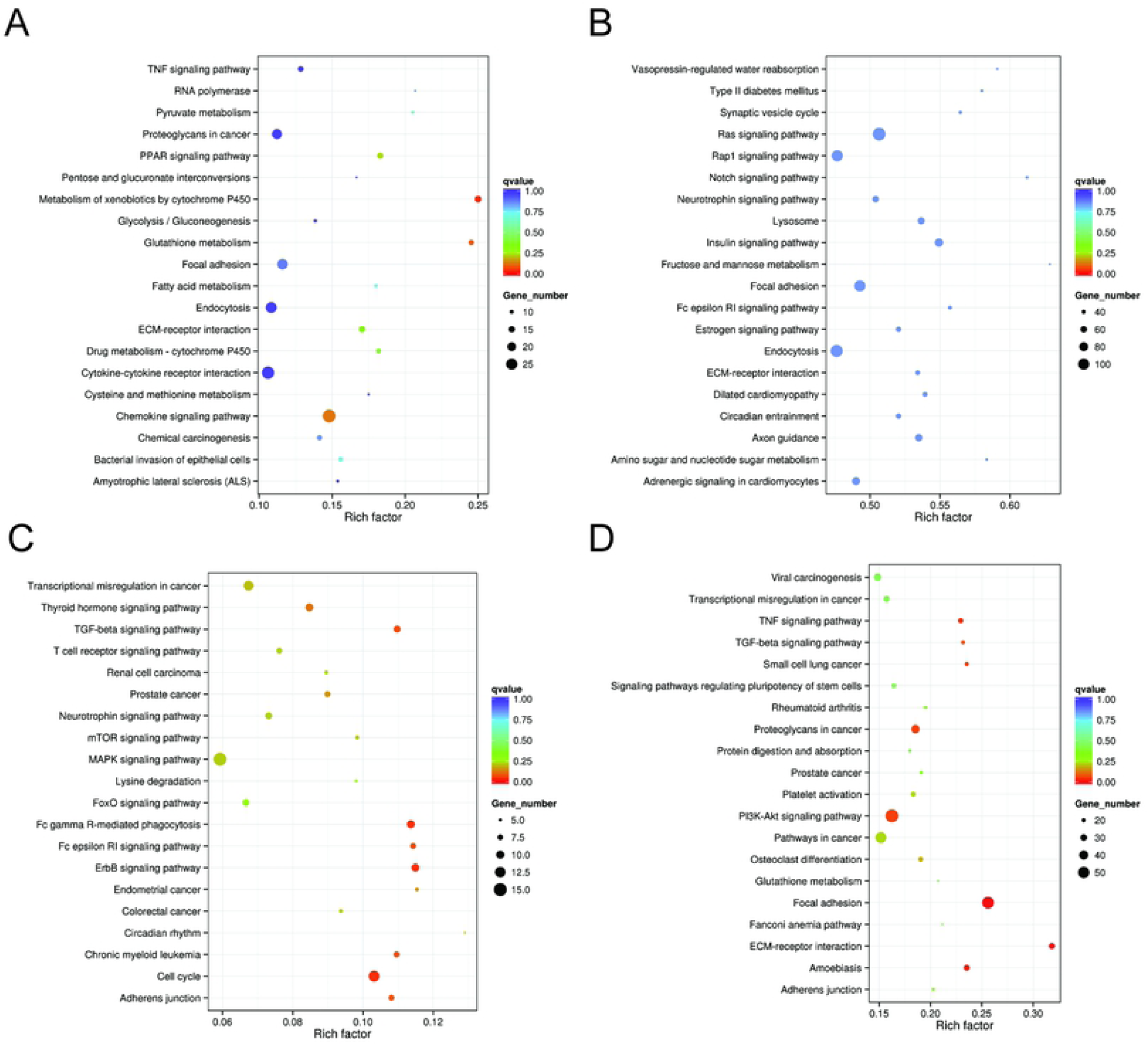
GC-1spg cell signaling pathways induced by BMP9 [A] The top 20 KEGG pathways with differentially expressed lncRNA, [B] miRNA, [C] circRNA, and [D] mRNA are shown.

## Discussion

BMP9, which belongs to the TGF-β superfamily, can induce MSCs, such as mouse embryonic fibroblasts (MEFs) or C3H10 cells, to differentiate into bone tissue [17,22,23,71,72]. Further, MSC-to-osteoblast transition is driven by BMP9, the most potent osteogenesis-inducing BMP signal *in vitro* and *in vivo* [21,22,73]. MSCs originating from the mesoderm can be isolated from different tissues [5,74,75,76,77]. Indeed, pluripotent stem cells isolated from testis are similar to MSCs, as demonstrated by gene expression profiling studies [5,77]. MSCs can be derived from the testis [5,8,74]. Though it is possible to differentiate bone tissue from MSCs, few studies have examined BMP9-induced osteogenesis of testis-derived MSCs.

GC-1spg cells, a mouse spermatogonia-derived cell line, represent the differentiation state of type B spermatogonia or preleptotene spermatocytes [1,37]. Several studies have demonstrated GC-1spg cell plasticity [1,4,78,79]. In our study, we drove BMP9 expression using a recombinant adenovirus. Real-time PCR results confirmed that our adenovirus stably overexpresses BMP9 in GC-1spg cells. Secondly, expression of osteogenic markers, including OPN, OCN, Sox9, and Runx2 [17,38,62,65,71,80], is enhanced by BMP9 in GC-1spg cells. Furthermore, ALP and mineralized calcium nodules, early osteogenesis markers, are increased by BMP9 overexpression. Therefore, our *in vitro* results demonstrate that exogenous BMP9 expression promotes osteogenic differentiation and mineralization in GC-1spg cells. To verify our *in vitro* results, we implanted BMP9-overexpressing cells in mice to induce ectopic ossification. Bony mass formation is augmented by exogenous BMP9 expression. Masson’s trichrome staining also showed that BMP9 increased trabeculae thickness, and alcian blue staining demonstrated that elevated BMP9 intensified cartilaginous matrix accumulation. Combined with Sox9 expression, we posit that chondrogenic signaling is important for BMP9-induced terminal osteogenic differentiation, although the underlying BMP9-mediated mechanism has not been thoroughly investigated.

BMP9 is one of the most potent osteogenic BMPs both *in vitro* and *in vivo* [21,22,25], playing a role in several diseases [29,35,69,81,82,83]. Signaling by BMP9 is a synergistic factor regulating the proliferation and migration of specific cell types and stem cell differentiation through extensive crosstalk with several signaling pathways. Though numerous signaling pathways are involved in BMP9-mediated cellular differentiation and proliferation, the cellular transduction mechanism of BMP9 remains unclear. To identify the signaling pathways activated by BMP9, we analyzed total RNA, including lncRNA, mRNA, circRNA, and miRNA, by RNA-seq and KEGG pathway analysis. Mechanistically, the TGF-β type I receptors ALK1 and ALK2 are required for BMP9-induced differentiation [67]. These receptors mediate nuclear signaling via Smad phosphorylation, suggesting that Smad or TGF-β signaling pathways mediate BMP9-induced osteogenesis in GC-1spg cells. Related pathways, including TNF [84,85], MAPK [86,87], Wnt [68], and Notch signaling pathways [14,27,53], may also play a role. In our study, KEGG pathway analysis of differentially expressed lncRNA, mRNA, circRNA, and miRNA shows that BMP9 treatment is involved in the cell cycle, focal adhesion, chemokine signaling, and the ErbB signaling pathway. These pathways are related to the MAPK signaling pathway [88], differentiation and angiogenesis [89], and the PPAR signaling pathway, which is involved in lipid metabolism and adipocyte differentiation [90].

Certain limitations have been identified in our study. MEFs and C3H10 are frequently used in osteogenesis studies, however, GC-1spg cells are rarely used in BMP9 research. This may explain why our results differ from previous findings on BMP9 activity. Indeed, BMP9-related signaling pathways were identified in MEFs, but not in GC-1spg cells. Secondly, we used adenovirus transfection to regulate BMP9 expression. It is possible that the signaling pathways identified in our study are induced by the adenovirus and not by BMP9 itself.

Collectively, our results indicate that the potent osteogenic signal BMP9 induces GC-1spg cells toward osteogenic differentiation *in vitro* and *in vivo*. Furthermore, high Sox9 expression and chondroid matrix accumulation in bony masses suggest that chondrogenesis might play a role in BMP9-induced bone formation. However, many signaling pathways other than MAPK and Notch are involved in BMP9-stimulated GC-1spg cells. Future studies investigating BMP9-mediated bone formation are warranted.

